# NSC-derived exosomes enhance therapeutic effects of NSC transplantation on cerebral ischemia in mice

**DOI:** 10.1101/2022.11.08.515655

**Authors:** Ruolin Zhang, Weibing Mao, Lumeng Niu, Wendai Bao, Yiqi wang, Zhihao Yang, Yasha Zhu, Haikun Song, Jincao Chen, Guangqiang Li, Meng Cai, Zilong Yuan, Jiawen Dong, Min Zhang, Nanxiang Xiong, Jun Wei, Zhiqiang Dong

## Abstract

Transplantation of neural stem cells (NSCs) has been proved to promote functional rehabilitation of brain lesions including ischemic stroke. However, the therapeutic effects of NSC transplantation is limited by the low survival and differentiation rates of NSCs due to the harsh environment in the brain after ischemic stroke. Here, we employed NSCs derived from human induced pluripotent stem cells (iPSCs) together with exosomes extracted from NSCs to treat cerebral ischemia induced by middle cerebral artery occlusion/reperfusion (MCAO/R) in mice. The results showed that NSC-derived exosomes significantly reduced the inflammatory response, alleviated oxidative stress after NSC transplantation, and facilitated NSCs differentiation *in vivo*. The combination of NSCs with exosomes ameliorated the injury of brain tissue including cerebral infarct, neuronal death and glial scarring, and promoted the motor function recovery. To explore the underlying mechanisms, we analyzed the miRNA profiles of NSC-derived exosomes and the potential downstream genes. Our study provided the rationale for the clinical application of NSC-derived exosomes as a supportive adjuvant for NSC transplantation after stroke.

## INTRODUCTION

Stroke is the second leading cause of death worldwide, which usually causes motor and cognitive impairments that require long-term rehabilitation (2021). The commonly used treatments of stroke in clinic include tissue plasminogen activator (t-PA) thrombolytic therapy and thrombus clearance surgery, but both are limited by inability to repair damaged neural circuits, and only 10% of stroke patients meet the treatment standards (Fugate and Rabinstein 2015, Nagaraja *et al.,* 2020). Stem-cell based therapy is a progressing and promising method to treat ischemic stroke. Many stem cell types including neural stem cells (NSCs) (Sakata *et al.,* 2012), embryonic stem cells (ESCs) (Meamar *et al.,* 2013), mesenchymal stem cells (MSCs) (Toyoshima *et al.,* 2017), bone marrow mononuclear cells (BMMCs) (Yang *et al.,* 2012), and iPSCs (induced pluripotent stem cells) (Duan *et al.,* 2021) have been tested in preclinical and clinical research, which showed encouraging therapeutic effects. Both endogenous and exogenous NSCs have remarkable capacity to maintain self-renewal while differentiating into various cell types including neurons and glial cells in nervous system (Dong *et al.,* 2012). iPSCs can be an ideal resource to acquire NSCs, which voids both ethical problems and immune rejection, and has a potential to provide genetically identical “patient-specific” cells for stroke patients (Baker *et al.,* 2019). On the other hand, the low survival rate of transplanted NSCs, largely due to chronic inflammation and oxidative stress of the microenvironment after stroke (Koutsaliaris *et al.,* 2022), and the poor differentiation of NSCs (Zhang *et al.,* 2019) limited its application.

NSC-derived exosomes are enriched in specific miRNAs that mediate multiple functions in physiological and pathological conditions (Luo *et al.,* 2022). NSC-derived exosomes, and have been proven useful for treating multiple neurological diseases due to their anti-inflammatory, neurogenic and neurotrophic effects as well as the interaction with the microenvironment of the brain tissue (Vogel *et al.,* 2018). Previous studies suggested that application of NSC-derived exosomes could promote the differentiation of NSCs through miRNAs *in vitro* (Yuan *et al.,* 2021). However, the effect of exosomes on grafted NSCs *in vivo* remains elusive. We propose that the combined treatment of exosomes and NSCs can effectively ameliorate harsh lesion conditions to help the NSCs survival and differentiation, achieving optimal treatment effects.

In this study, we established ischemic stroke in mice with MCAO/R, and tested different treatment strategies using transplantation of iPSC-induced NSCs and NSC- derived exosomes. Our results indicated that NSC-derived exosomes could promote NSCs differentiation, reduce the oxidative stress and inflammation, and alleviate the formation of glial scars after ischemia and reperfusion, and as a result, could enhance the therapeutic effects of NSC transplantation. We further explored the molecular mechanisms through profiling the miRNAs of the NSC-derived exosomes.

## RESULTS

### NSC-derived exosomes facilitated post-stroke recovery after NSC transplantation in MCAO/R mice

We first characterized the NSCs derived from iPSCs by examining the expression of NSC marker genes including *SOX2* and *PAX6* by immunocytochemistry staining. The results showed that the cells used for subsequent transplantation expressed high level of NSC marker genes (Figure 1 - supplement 1A), indicating that NSCs were efficiently induced from iPSCs. We isolated exosomes from the same NSCs and examined the expression of exosomal markers including TSG101, CD63, and CD9 (Figure 1 - supplement 1B). Furthermore, the results of transmission electron microscopy (TEM) showed that the particle size of exosomes mixture was less than 200 nm, and Nanoparticle Tracking Analysis (NTA) confirmed the typical distribution of particle diameter of exosomes (Figure 1 - supplement 1C and D).

**Figure 1.**
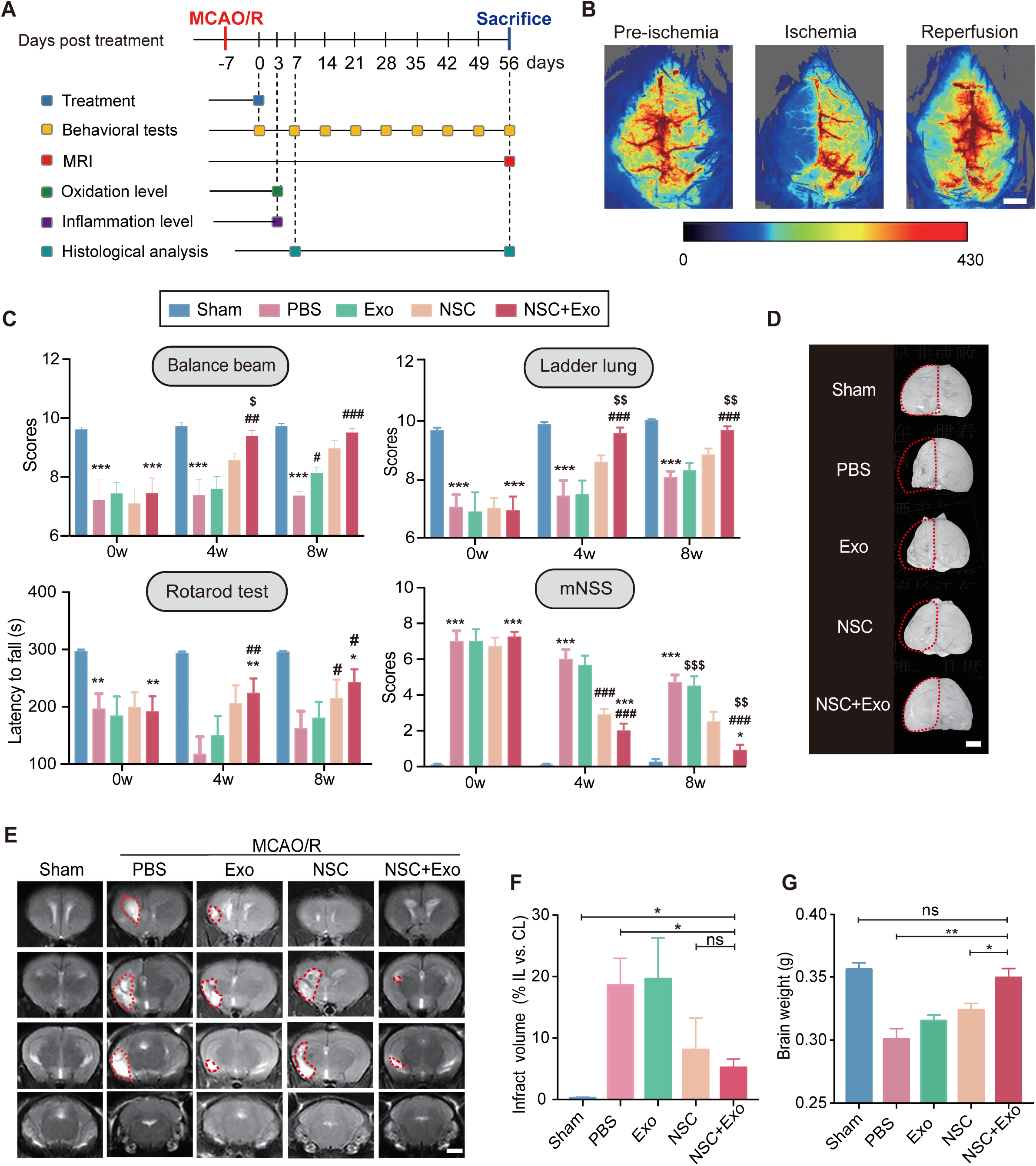
NSC-derived exosomes enhanced the therapeutic effects of NSCs on motor impairment and brain infarction after stroke. (A) Summary of the experimental timeframes. (B) Images of cerebral blood flow before, during and 24 h of the MCAO/R procedure. Scale bar: 2 mm. (C) Behavioral test results (the balance beam, ladder rung, rotarod tests) and mNSS at 0, 4 and 8 weeks after treatment, n = 10 mice per group. \**P* < 0.05, \*\**P* < 0.01, \*\*\**P* < 0.001, versus Sham group. ^#^*P* < 0.05, ^##^*P* < 0.01, ^###^*P* < 0.001, versus PBS group. ^$^*P* < 0.05, ^$$^*P* < 0.01, ^$$$^*P* < 0.001, versus NSC group. (D) Representative images show brain atrophy at 8 weeks after treatment. The ischemic hemispheres are marked by dotted lines. Scale bar: 2 mm. (E) MRI images show brain cerebral infarct at 8 weeks after treatment. The infarct area is marked by dotted lines. Scale bar: 2 mm. (F) Quantification of (E), n = 3 per group. (G) Quantification of brain weights at 8 weeks after treatment. \**P* < 0.05, \*\**P* < 0.01, \*\*\**P* < 0.001, ns indicates non-significant difference. Figure 1 **- source data 1.** NSC-derived exosomes enhanced the therapeutic effects of NSCs on motor impairment and brain infarction after stroke.

We next examined the effects of different treatment strategies on the brain lesion after cerebral ischemia and reperfusion in MCAO/R mice. To examine the presence and persistence of cerebral edema, TTC staining was performed 1 and 7 days after MCAO/R (Figure 1 - supplement 1E). Two doses of NSC transplantation, 2×10^5^ and 5×10^5^, were first tested and the results of survival curve (Figure 1 - supplement 1F) and rotarod test (Figure 1 - supplement 1G) showed a dose-dependent effects of transplanted NSCs. Therefore, we determined to use the dose of 5×10^5^ NSCs for the subsequent treatments to achieve a robust therapeutic effect. Mice were randomly divided into five groups (Sham, PBS, Exo, NSC and NSC+Exo). Except Sham group, mice in all the other four groups received standard MCAO/R surgery. Lateral ventricle injections of 5 µL PBS (PBS group), 10 µg exosomes in 5 µL PBS (Exo group), 5 × 10^5^ NSCs in 5 µL PBS (NSC group), and 5 × 10^5^ NSCs + 10 µg exosomes in 5 µL PBS (NSC+Exo group) were performed at 7 days post-MCAO/R (Figure 1A). The levels of reactive oxygen species (ROS) and inflammation were measured in focal brain tissues at 3 days post treatment; behavioral assessments were performed at 0 to 8 weeks post treatment; histological examinations were analyzed at 8 weeks post treatment (Figure 1A). To ensure the successful establishment of cerebral ischemia, the cerebral blood flow was examined before, during and after MCAO/R (Figure 1B). Neurological functions were evaluated by balance beam, ladder lung, rotarod test and Modified Neurological Severity Score (mNSS) up to 8 weeks after treatment (Figure 1C and Figure 1 - supplement 2A-D). The results suggested that transplantation of NSCs combined with exosomes began to take effect starting at 4 weeks after treatment, as evidenced by the results of balance beam, ladder lung and mNSS test, and significantly worked better than that solely with NSCs at 8 weeks post treatment (Figure 1C). The infarct area in the ipsilateral hemisphere was determined by MRI (Figure 1E) at 8 weeks post treatment. Compared to the severe damages of brain tissues in PBS group, mice treated by NSCs combine with exosomes showed significantly reduced infarct areas (Figure 1F). Meanwhile, the combination of NSCs and exosomes showed better protective effects on the brain tissue than either alone (Figure 1D), which was further confirmed by the results of brain weight analysis (Figure 1G). Therefore, our results indicated that NSC-derived exosomes could significantly enhance the therapeutic effects of NSCs on motor dysfunction and brain infarction in MCAO/R mice. Furthermore, the NSCs-mediated therapeutic effects were greatly accelerated by addition of exosomes.

### NSC-derived exosomes enhanced the therapeutic effects of NSCs on neuronal damage

We next examined the recovery of ischemia-induced neuronal damage of cerebral cortex in different treatment groups. The results of NeuN staining revealed that, compared to the mice treated solely with NSCs, the combination NSCs and exosomes significantly reduced the tissue loss from 14.32±3.52% to 7.57±2.59% (Figure 2A, B), while administration of exosomes alone showed no significant difference compared to PBS treatment (Figure 2B). The reduction of infarct area after combined treatment was also accompanied with improved dendritic density and length (Figure 2C, D), alleviated spines loss (Figure 2E), and increased complexity of neuronal projections (Figure 2F) in the cerebral cortex. Interestingly, although exosome treatment did not show robust therapeutic effects on behavior impairment and infarct area, the number of dendritic spines was significantly increased by exosome treatment in Exo group compared to that of PBS group (Figure 2E), suggesting that exosomes might play an important role in the recovery of neuronal complexity. MCAO/R mice had damaged pyramidal and granular cells with fuzzy cell contours as shown by Nissl staining (Figure 2G and Figure 2 - supplement 1). The addition of exosomes could further reduce the neuronal loss in the ipsilesional hemisphere on top of the effects of NSC transplantation.

**Figure 2.**
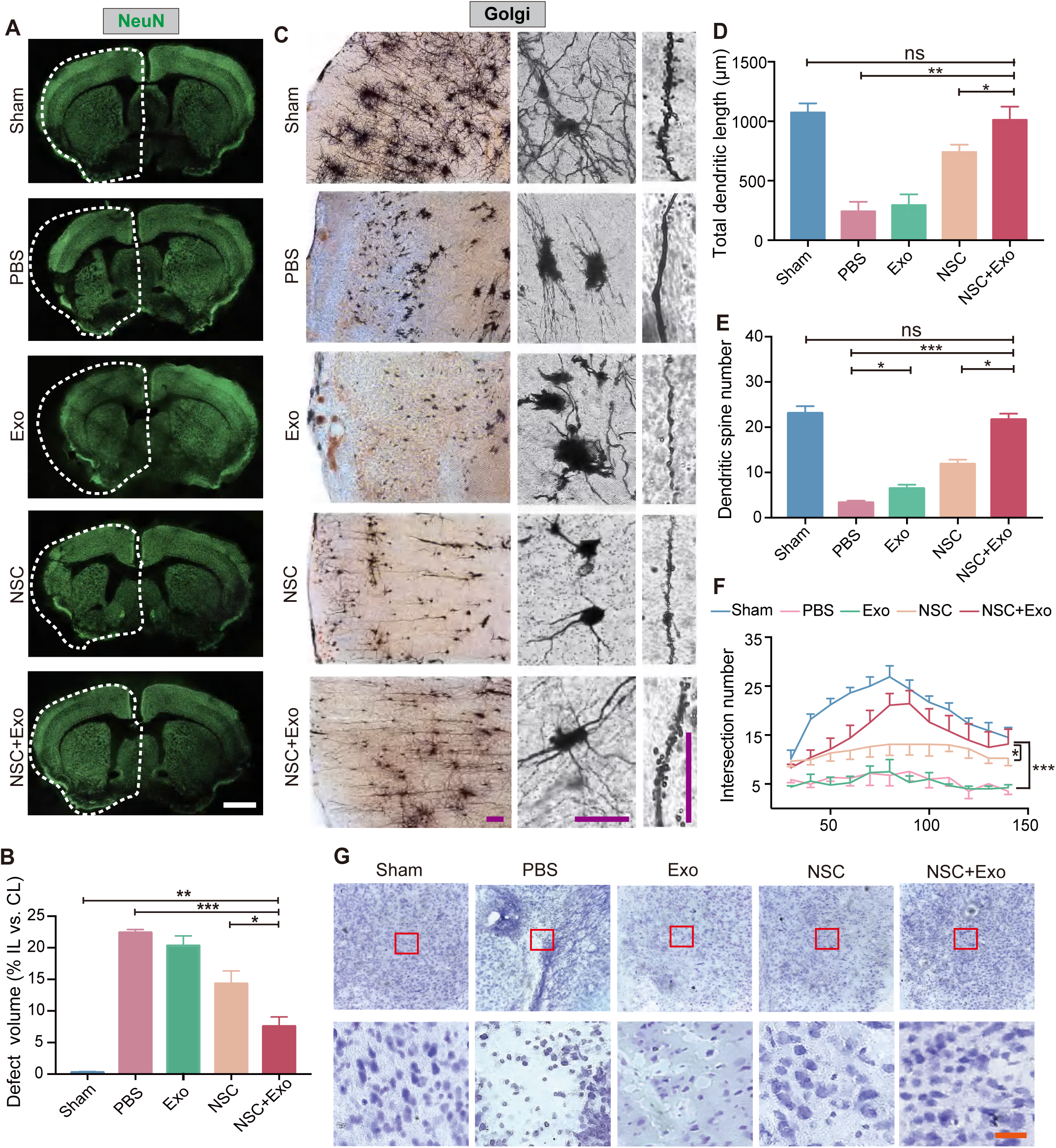
Effects of combined treatment with NSCs and exosomes on neuronal damage in MCAO/R mice. (A) Immunofluorescent staining of NeuN of different groups at 8 weeks after treatment. The ipsilateral hemispheres were marked by the dotted lines. Scale bar: 2mm. (B) Quantification of defect volume of NeuN staining, n = 4 mice per group. (C) Representative images of Golgi-Cox staining in the infarct area at 8 weeks after treatment. Quantitative analysis of total dendritic length (D), dendritic spine number (E), and neuronal complexity (F). Scale bar: 100 µm. Fifteen neurons from 3 mice were analyzed for each group. (G) Nissl styining of infarct area in the brain at 8 weeks after treatment. Scale bar: 25 µm. \**P* < 0.05, \*\**P* < 0.01, \*\*\**P* < 0.001. ns denotes not significant. **Figure 2 - source data 1.** Effects of combined treatment with NSCs and exosomes on neuronal damage in MCAO/R mice.

### Exosomes promoted the differentiation of transplanted NSCs

To investigate the regulatory effects of NSC-derived exosomes on transplanted NSCs, the differentiation rates of NSCs in the cerebral cortex *in vivo* were evaluated at 8 weeks after transplantation. Compared to the NSC group, the number of tdTomato positive NSCs was significantly increased in the NSC+Exo combined treatment group (Figure 3A, C). Among the tdTomato positive cells, Nestin^+^/tdTomato^+^ cells were less in NSC+Exo group than the other groups (Figure 3A, D), while the number of Tuj1^+^/tdTomato^+^ cells was significantly higher in NSC+Exo group, which implied that exosomes could promote the differentiation of NSCs into neurons (Figure 3B, E). The TUNEL staining indicated that the excessive apoptosis cells after MCAO/R were also reduced by exosome transplantation (Figure 3 - supplement 1). Therefore, our data indicated that co-transplantation of exosomes could effectively facilitate the differentiation of transplanted NSCs in MCAO/R mice.

**Figure 3.**
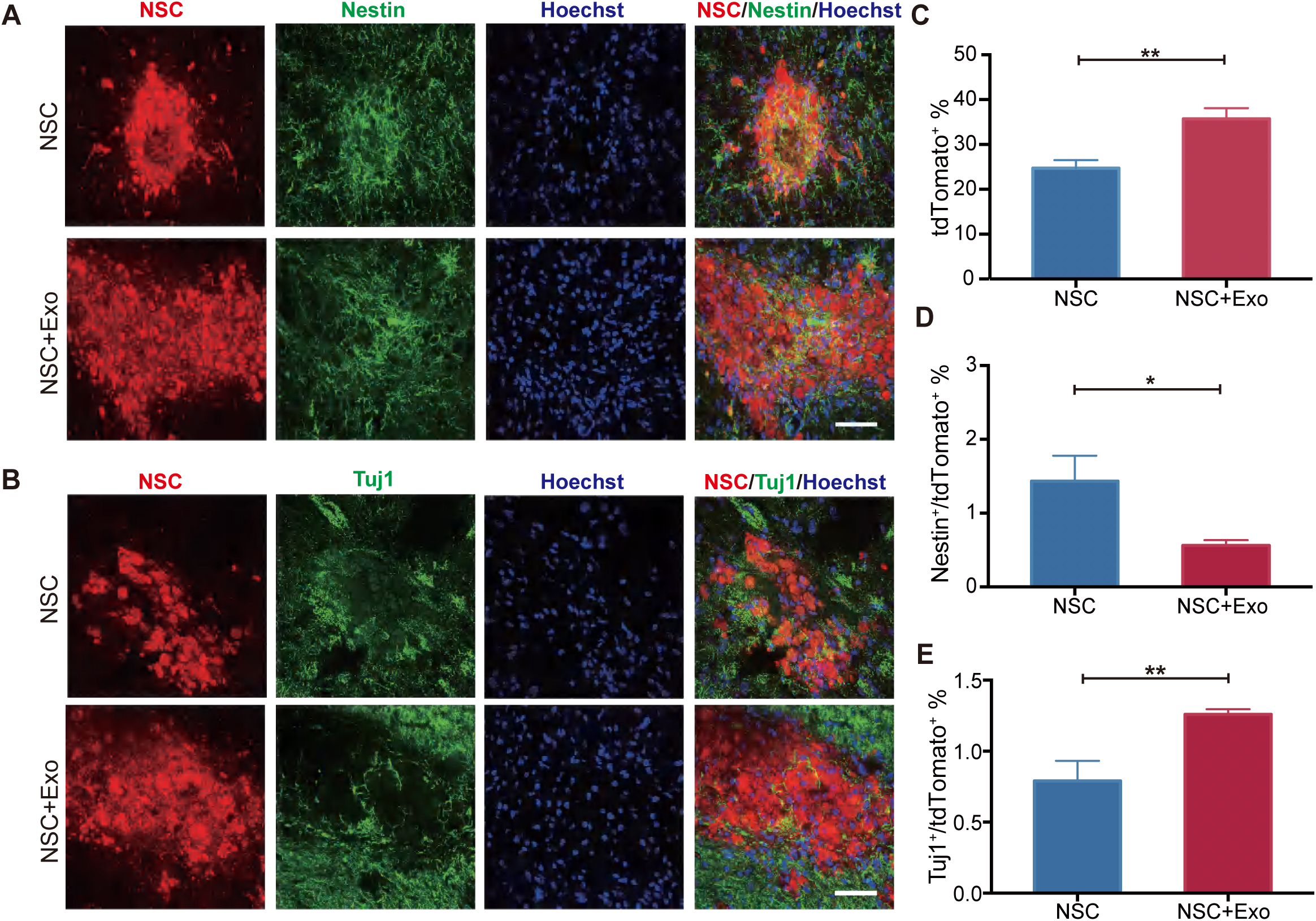
Effects of NSC-derived exosomes on the differentiation of transplanted NSCs. (A) Representative images of Nestin staining (green) at 8 weeks after transplantation. (B) Tuj1 staining (green) at 8 weeks after transplantation. Nuclei are counterstained with Hoechst (blue in A and B). Quantification of tdTomato^+^ cells (C), Nestin^+^/tdTomato^+^ cells (D), and Tuj1^+^/ tdTomato^+^cells (E). Scale bar: 50 µm. \**P* < 0.05, \*\**P* < 0.01. Figure 3 **- source data 1.** NSC-derived exosomes promoted the differentiation of transplanted NSCs.

### Exosomes promoted the microenvironment remodeling

Oxidative stress and global brain inflammation are closely involved in the progressing pathology after stroke (Hurn *et al.,* 2007, Shi *et al.,* 2019), which challenges the survival and colonization of transplanted NSCs (Li *et al.,* 2017). We employed oxygen and glucose deprivation (OGD)/re-oxygenation (OGD/R) procedure on cultured NSCs to simulate the main pathogenesis of stroke, ischemia-reperfusion (Zhang *et al.,* 2017, Yu *et al.,* 2018). The results showed that OGD/R treatment could induce high level of oxidative stress in NSCs, whereas exosomes could reduce the production of ROS after OGD/R (Figure 4A and 4B). We further examined the expression of oxidative stress-related genes. The mRNA expression level of *CHOP* (endoplasmic reticulum stress marker) was reduced by exosome treatment after OGD/R (Figure 4C). Meanwhile, exosome treatment increased the expression of antioxidant genes *NRF2, NQO1* and *SOD2* (Figure 4C). Besides the *in vitro* OGD/R experiments, the level of oxidative stress *in vivo* was also determined at 3 days post stroke, and the results showed that the MDA content was significantly decreased in exosome-treated mice (Figure 4D). Therefore, our data suggested that NSCs-derived exosomes could ameliorate oxidative stress, which could potentially facilitate the survival, colonization and differentiation of transplanted NSCs.

**Figure 4.**
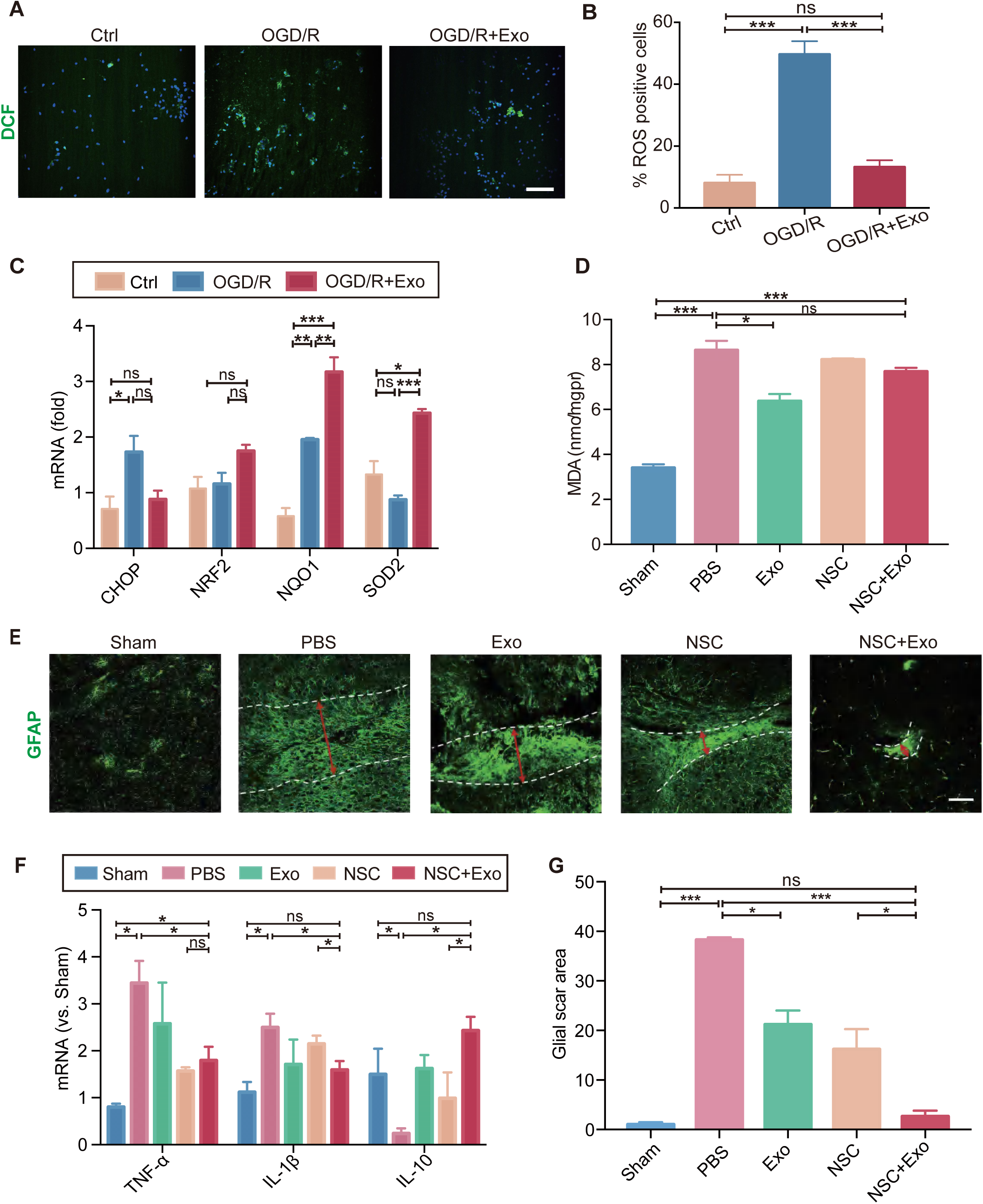
Effects of exosomes on the microenvironment remodeling. (A) ROS generation was evaluated by DCF-DA fluorescent probe labeling (green) in OGD/R-treated NSCs. Nuclei were counterstained with Hoechst (blue). Scale bar: 50 µm. (B) Percentage of ROS positive cells. (C) The relative mRNA expression of oxidative stress-related genes *CHOP*, *NRF2, NQO1* and *SOD2* were measured by RT-qPCR. (D) The MDA level at 3 days after treatment, n = 3 mice / group. (E) Representative images and quantification (G) of scar-forming astrocytes detected by GFAP staining. Scale bar: 50 µm. (F) The mRNA expression of *TNF-α*, *IL-1β* and *IL- 10* of the ipsilesional brain was measured by RT-qPCR at 3 days after treatment, n = 3 mice / group. \**P* < 0.05, \*\**P* < 0.01, \*\*\**P* < 0.001. ns indicates non-significant difference. Figure 4 **- source data 1.** Effects of exosomes on the microenvironment remodeling.

Heterologous stem cells transplantation could induce robust inflammatory response. It has been reported that the proliferation of immune cells reaches the peak during the acute phase post-transplantation (Graf and Stern 2012, Boncoraglio *et al.,* 2019). Interestingly, our results showed that exosomes could reduce the expression of inflammatory cytokines including *TNF-α* and *IL-1β,* while increase the expression of anti-inflammatory cytokine *IL-10* in NSCs after OGD/R (Figure 4F) suggesting that exosomes could alleviate the elevated immune response after NSC transplantation.

After brain tissue damages caused by conditions such as cerebral ischemia and reperfusion, glial scars are usually formed, which can reestablish the physical and chemical integrity of the brain tissue by generating a barrier across the injured area, but inhibit the neuronal recovery as well (Michinaga and Koyama 2021). Astrocytes are the main cellular component of the glial scar, and our results indicated that astrocytes were prone to form glial scars during the chronic phase after stroke (Figure 4E and Figure 4 - supplement 1A). We subsequently investigated the effects of different treatments on the formation of glial scars in MCAO/R mice. The results suggested that the combined treatment of NSCs and exosomes significantly decreased the glia scars in the subacute phase (Figure 4 - supplement 1B) and the chronic phase (Figure 4E and 4G).

### miRNA profiling and functional enrichment analysis of NSC-derived exosomes

To explore the underlying molecular mechanisms of exosomes regulating the transplanted NSCs and the microenvironment, we proposed that the exosomes might regulate target genes through the release of miRNAs, components of the key functional molecules carried by exosomes. Therefore, we profiled the miRNA expression of NSC- derived exosomes using miRNA microarray. A total of 850 known miRNAs were detected, and the top 10 miRNAs with the highest read counts were displayed and verified by qPCR (Figure 5A and Figure 5 - supplement 1A). Targetscan, miRcode and miRDB databases were used to predict the downstream targets of the top 10 abundant miRNAs, and 17 potential relative target genes were selected, which have been proved to play important roles in neural modulation. The interactive network of the exosomal miRNAs and the selected target genes were analyzed and visualized using Cytoscape (Figure 5B). Target genes were predicted to be regulated by multiple miRNAs, among which hsa-miR-30a-5p and hsa-miR-7-5p were involved in multiple regulation. We next examined the effects of exosome treatment on the expression of candidate target genes in NSCs after OGD/R by RT-qPCR using *STAT3*, *PTPN1* and *CHUK* as examples (Figure 5 - supplement 1B) (Park *et al.,* 2012, Wang *et al.,* 2018, Culley *et al.,* 2019). The results showed that exosomes reduced the expression of downstream genes in NSCs, which was consistent with our hypothesis that exosomes regulate the recipient cells through carrying miRNAs that downregulate the target genes.

**Figure 5.**
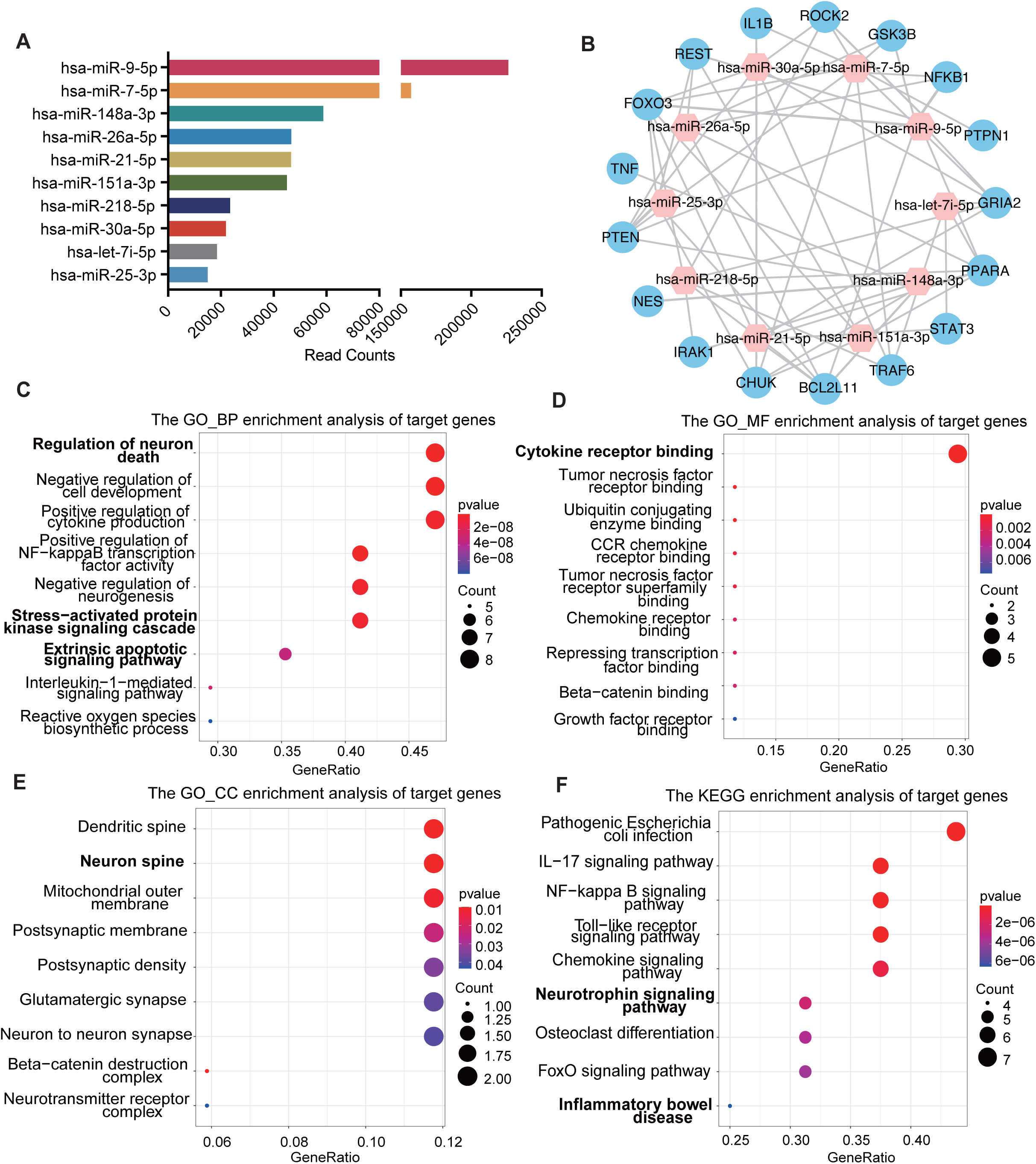
miRNA profiling of NSC-derived exosomes and enrichment analysis of predicted target genes. (A) The read counts of the top 10 abundant miRNAs in NSC-derived exosomes. (B) miRNA-mRNA regulatory networks. miRNA and mRNA are represented by the pink and blue circles respectively. Gene ontology analysis of predicted target genes in terms of biological process (C), molecular function (D) and cellular component (E). (F) The KEGG pathway analysis of predicted target genes.

To further understand the regulatory effects of exosomal miRNAs on NSCs and the microenvironment, we performed GO enrichment analysis and KEGG pathway analysis on all the potential target genes. GO enrichment analysis, in terms of biological process (BP, Figure 5C), molecular function (MF, Figure 5D) and cellular component (CC, Figure 5E), disclosed that the potential target genes were enriched in functions that were correlated with cellular and microenvironmental homeostasis of the central nervous system such as regulation of neuron death and neurogenesis, stress−activated protein kinase signaling cascade, cytokine receptor binding, and neuron spine. KEGG pathway analysis suggested that the target genes were mainly involved in inflammation and apoptosis-related signaling pathways (Figure 5F). Therefore, the predicted target genes of exosomal miRNAs were concentrated in the functions and pathways that could regulate the cellular behavior of transplanted NSCs as well as the microenvironment remodeling.

Taken together, our findings suggested that NSC-derived exosomes might regulate the transplanted NSCs and the surrounding microenvironment through carrying the miRNAs which could further modulate the downstream genes and pathways in both the NSCs and the surrounding cells (Figure 6).

**Figure 6.**
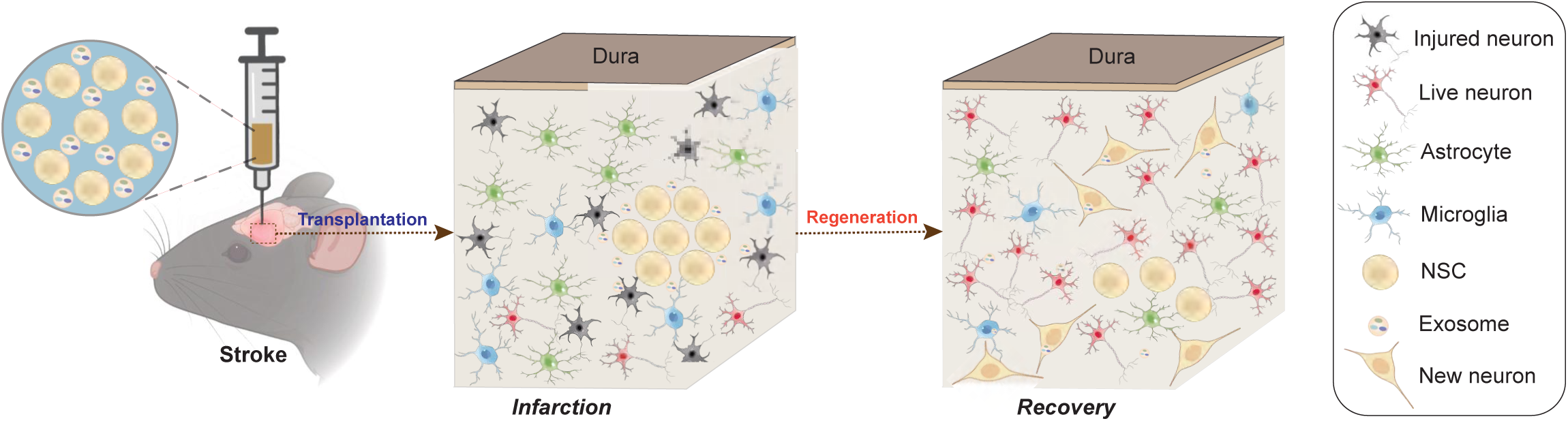
Schematic illustration of the mechanisms for combination treatment of NSCs and exosomes in neuroprotection against ischemic stroke. NSCs and exosomes achieve therapeutic goals by repairing damaged neurons, alleviating the inflammatory environment and reducing glial cell activation.

## DISCUSSION

Stem cell-based therapy is an emerging and promising method to treat stroke, due to its effectiveness in cell replacement, neuroprotection, angiogenesis, and modulation on inflammation and immune response (Hao *et al.,* 2014a), but poor survival and differentiation of grafted cells have limited its efficacy and application (Jiang *et al.,* 2019). In the present study, we co-transplanted NSCs-derived exosomes with NSCs in MCAO/R -induced cerebral ischemia in mice. Consistent with previous reports (Wei *et al.,* 2017, Zhang *et al.,* 2019), our results confirmed that NSCs could effectively promote the recovery of motor function post stroke in mice. Importantly, we demonstrated that exosomes could promote the repairment of the damaged brain tissue as well as the functional recovery, enhance the differentiation of the grafted NSCs in the infarct area, reduce the oxidative stress and inflammation, and alleviate the formation of glial scars in MCAO/R mice. As a proof-of-concept study on the co-delivery of NSCs together with exosomes in the classical animal model of ischemic stroke, our study provided solid rationale supporting the application of exosomes during stem cell-based therapy. On the other hand, whether exosomes from different sources have similar effects on transplanted NSCs and how exosomes regulate other types of stem cells *in vivo* deem further exploration.

The pathological process of ischemia-reperfusion includes the generation of ROS, brain edema, and the increased levels of inflammation, which leads to the tough microenvironment for the transplanted stem cells to survive and differentiate (Zhang *et al.,* 2019). In addition, transplanted allogeneic stem cells also exacerbate oxidative stress levels. Yahata, T *et al* found that transplantation of human hematopoietic stem cells triggers replication stress and induces increasing ROS levels in mice (Yahata *et al.,* 2011). Bone marrow mesenchymal stem cells (BMSCs) transplantation has also been reported to increase oxidative stress levels in mouse muscle cells (Liu *et al.,* 2019). Besides the oxidative stress, the chemokines released by macrophages and endothelial cells after stroke, such as chemokine group CXC ligand 1 (CXCL1), recruit peripheral immune cells to flood into the damaged brain, which causes immune-inflammatory damage (Ormstad *et al.,* 2011). Meanwhile, transplantation of exogenous stem cells could aggravate the inflammatory response around infract area due to the immune-rejection (Hao *et al.,* 2014b). Xia *et al* reported that ESC-derived exosomes decrease the inflammatory response, alleviate neuronal death, and improve long-term recovery after MCAO/R through increasing regulatory T cells (Xia *et al.,* 2021). Due to the properties of regulating signaling pathways in target cells, and remodeling the microenvironment (Vogel *et al.,* 2018), NSCs-derived exosomes have been demonstrated to improve a variety of neurological diseases, such as Alzheimer’s disease (Liu *et al.,* 2020), spinal cord injury (Ma *et al.,* 2019), and ischemic stroke (Sun *et al.,* 2019). Recent evidence demonstrated that exosomes promote the maturation of both neuron and glial cells *in vitro* (Yuan *et al.,* 2021). Furthermore, excessive initiation of apoptosis has also been implicated in stroke (Hwang *et al.,* 2013). Here we showed that NSC-derived exosomes could reduce the oxidative stress and the inflammatory response, and promote the differentiation of transplanted NSCs and reduce excessive apoptosis in MCAO/R mice. Therefore, our results indicated that exosomes could promote the therapeutic effects of transplanted NSCs at multiple levels.

Previous studies have shown that stem cell-derived exosomes had neural protective effects and could promote recovery after ischemic stroke (Webb *et al.,* 2018, Sun *et al.,* 2019, Xia *et al.,* 2021). However, we did not observe significant therapeutic effects with solely exosome treatment, which could be due to the dose of exosomes, the treatment timing and frequency. Considering the fact that cell transplantation requires a relatively stable microenvironment (Nih *et al.,* 2017, Lee *et al.,* 2018), we transplanted NSCs and exosomes at 7 days after stroke without subsequent delivery of exosomes in this study. Although the delivery of exosomes alone used in this study did not show significant neural protective effects, it indeed ameliorated oxidative and inflammatory lesion conditions, promoted neuronal repairment, and potentiated the therapeutic power of transplanted NSCs, suggesting that the application of exosomes could be an effective adjuvant for NSC-based therapy. Besides, as exosomes are ideal carriers for drug delivery (Chen *et al.,* 2021), modifications of exosomes by adding drugs or other functional molecules could potentially further enhance the beneficial effects of exosome treatment.

As miRNAs were reported to be one of the major exosomal components, we profiled the miRNAs from NSCs-derived exosomes to explore the molecular basis for the effects of exosomes as we observed in this study. Bioinformatic enrichment analysis in this study suggested that the predicted garget genes of exosomal miRNAs were concentrated in the functions and pathways that could regulate the NSCs’ behavior and the microenvironment. Interestingly, inflammation and oxidative stress-related genes and signaling pathways were also highly enriched in the target genes, consistent with the antioxidant role of exosomes as disclosed by our study. We, therefore, provided clues and a useful resource of exosomal miRNAs and predicted target genes for understanding the mechanisms underlying the function of exosomes in the NSC-based therapy for ischemic stroke. The roles and working model of the exosomal miRNAs as well as predicted target genes demand further exploration.

## MATERIALS AND METHODS

### Key resources table

| Reagent type (species) or resource | Designation | Source or reference | Identifiers | Additional information |
| --- | --- | --- | --- | --- |
| Cell line (Homo sapiens) | iPSCs | (Cai *et al.*, 2021) |  |  |
| Commercial assay or kit | BCA assay | Biosharp | BL521A |  |
| Chemical compound, drug | BSA | Sigma |  |  |
| Antibody | Anti -Sox2 | GeneTex | Cat. No 43019 | 1:500 |
| Antibody | Anti-E-Cadherin | Proteintech | Cat. No 60335 | 1:200 |
| Antibody | Anti-Pax6 | Proteintech | Cat. No 12323 | 1:200 |
| Antibody | Anti-Ki67 | Cell signaling technology | Cat. No 9449 | 1:1000 |
| Chemical compound, drug | Goat anti-Mouse IgG 568 | Invitrogen | A-10037 | 1:500 |
| Chemical compound, drug | Goat anti-Rabbit IgG 568 | Invitrogen | A-11036 | 1:500 |
| Chemical compound, drug | Goat anti-Mouse IgG 488 | Invitrogen | A-11029 | 1:500 |
| Chemical compound, drug | Goat anti-Rabbit IgG 488 | Invitrogen | A-11034 | 1:500 |
| Antibody | Anti-NeuN | Abcam | Cat. No ab177487 | 1:300 |
| Antibody | Anti-GFAP | Cell signaling technology | Cat. No 3670S | 1:1000 |
| Antibody | Anti-β-III-tubulin | Cell signaling technology | Cat. No 5568S | 1:300 |
| Antibody | Anti-Nestin | Santa Cruz Biotechnology | Cat. No Sc-23927 | 1:250 |
| Antibody | Anti-TSG101 | ABclonal | Cat. No A1692 | 1:1000 |
| Antibody | Anti-CD9 | ABclonal | Cat. No A19027 | 1:2000 |
| Antibody | Anti-CD63 | ABclonal | Cat. No A19023 | 1:1000 |
| Chemical compound, drug | Goat anti-Mouse | Proteintech | Cat No. PR30012 | 1:5000 |
| Chemical compound, drug | Goat anti-Rabbit | Proteintech | Cat No. PR30011 | 1:5000 |

| Reagent type<br>(species) or<br>resource | Designation | Source or<br>reference | Identifiers | Additional<br>information |
| --- | --- | --- | --- | --- |
| Commercial<br>assay<br>or kit | ECL Enhanced<br>Kit | Abclonal | RM00021 |  |
| Chemical<br>compound,<br>drug | Isoflurane | RWD | R510-22-10 |  |
| Chemical<br>compound,<br>drug | RIPA lysis<br>buffer | Biosharp | BL504A |  |
| Commercial<br>assay<br>or kit | FD rapid Golgi<br>Stain kit | FD Neuro<br>Technologies | PK401A |  |
| Software,<br>algorithm | Fiji | (Schindelin *et al.*,<br>2012) |  | https://imagej.net/Fiji<br>i |
| Commercial<br>assay<br>or kit | Nissl Stain<br>Solution | Solarbio | G1434 |  |
| Chemical<br>compound,<br>drug | DCF-DA | Invitrogen | D399 | 10 µM |
| Chemical<br>compound,<br>drug | 4%<br>paraformaldehy<br>de | Biosharp | BL539A |  |
| Chemical<br>compound,<br>drug | Triton X-100 | Biofroxx | Cat. No.<br>1139ML100 |  |
| Chemical<br>compound,<br>drug | 10% formalin | Coolaber | SL1560 |  |
| Chemical<br>compound, drug | OCT compound | SAKURA | Japan |  |
| Software,<br>algorithm | Targetscan |  |  | http://www.targetscan.org/ |
| Commercial<br>assay<br>or kit | Cell death<br>detection kit | Roche | Cat. No.<br>11684795910 |  |

| Reagent type<br>(species) or<br>resource | Designation | Source or<br>reference | Identifiers | Additional<br>information |
| --- | --- | --- | --- | --- |
| Commercial<br>assay<br>or kit | TBA method | Nanjing Jiancheng<br>Bioengineering<br>Institute | A003-1 |  |
| Software,<br>algorithm | miRcode |  |  | http://www.mircode.org/ |
| Software,<br>algorithm | miRDB |  |  | http://mirdb.org/ |
| Software,<br>algorithm | Cytoscape<br>software |  |  |  |
| Software,<br>algorithm | RStudio | RStudio |  |  |
| Software,<br>algorithm | GraphPad<br>PRISM | GraphPad<br>Software | Version 9.2.0 |  |

### Animals

Male C57BL/6 mice (age: 7 - 8 w, weight: 22 - 24 g) were selected due to estrogen and progesterone have recognized neuroprotective effects (Newton *et al.,* 2022). All animal procedures were performed in compliance with guidelines for the care and use of animals and were approved by the University Huazhong Agriculture Institutional Animal Care and Use Committee (Approval number: HZAUMO-2021-0111). Mice were assigned to MCAO/R or sham operation, and accepted NSCs or exosome treatment.

### NSCs induction and culture

NSCs induction was performed by iRegene Therapeutics, Wuhan, China as previously reported (Cai *et al.,* 2021). Briefly, Human iPSCs were cultured with STEMdiff Neural Induction Medium (Stemcell Technologies). The medium was replaced daily until day 9. After the first passage, Y-27632 was added to the medium on day 1 to ensure the cell attachment and then removed from the medium on day 2. NSCs were cultured in STEMdiff Neural Progenitor Medium (Stemcell Technologies) to maintain cell growth after passage. NSCs were identified by immunofluorescence staining and qPCR.

### Exosomes isolation and detection

Cellular debris were removed from cell culture supernatant at 2000 g for 10 min. The supernatants were centrifuged at 2,0000 g for 30 min. Then, exosomes were collected by ultracentrifugation (Beckman, America) at 100,000g for 120 min. Finally, exosomes were washed in 12 ml PBS and collected again for 90 min. Exosomes were resuspended in PBS and protein concentration was measured by BCA assay (Biosharp, China). For observation by transmission electron microscopy, exosomes were fixed in 2.5% glutaraldehyde at 4 °C overnight and then mounted on a copper grid, stained with 2% uranyl acetate, and examined with a transmission electron microscope (Japan) with 100 kV. For nanoparticle tracking analysis, exosomes were examined by Malvern Nano ZS90 as previously described (Shi *et al.,* 2018). Exosomes were diluted in PBS and 1.0 mL suspension was loaded into a cuvette to measure and analyze.

### MCAO/R model

Ischemic stroke was established with MCAO/R surgery on male C57BL/6 mice (age: 7-8weeks, weight: 22-24g). Mice were anesthetized with 2% isoflurane (RWD, China). For focal cerebral ischemia, a silicon-coated filament (RWD, China) was inserted into the left middle cerebral artery (MCA) to block blood flow. Sixty minutes later, the filament was extracted for reperfusion. Rectal temperature was maintained at 37°C during the entire procedure. Then anesthesia was discontinued and mice were allowed to recover. The cerebral blood flow (rCBF) was detected using a laser doppler flowmetry (Perimed, Sweden). A 55% decrease in the rCBF of the ipsilateral hemisphere, as compared to contralateral hemisphere, was considered the threshold for successful establishment of cerebral ischemia. Mice of the Sham group were performed the same as the MCAO/R procedure without filament insertion.

### Delivery of NSCs and exosomes

Eighty-eight MCAO/R mice were randomly divided into 4 groups at 7 days post operation, and twelve mice with low body weight (less than 15g) were excluded. Mice were anesthetized and placed in a mouse stereoscopic apparatus (RWD, China). The skull was drilled to make a burr hole above the lateral ventricle (AP+0, ML-1, DV-2.25 mm) for NSCs and exosomes injection. NSCs were genetically labeled with tdTomato for cell tracking. The five groups were treated as follows: Model (MCAO/R mice treated with 5 µL PBS), Exo (MCAO/R mice treated with 10 µg exosomes in 5 µL PBS), NSC (MCAO/R mice treated with 5×10^5^ NSCs in 5 µL PBS), NSC+Exo (MCAO/R mice treated with 5×10^5^ NSCs combine with 10 µg exosomes in 5 µL PBS) and Sham (fifteen mice with sham operation not treated).

### Immunofluorescence staining

The cells were planted on round glass coverslips, and fixed with 10% formalin overnight at 4 °C, then permeabilized with 0.25% Triton X-100, and blocked with 2% BSA (Sigma, America) for 1h at room temperature. The coverslips were incubated with primary antibodies including anti-Sox2 (1:500, GeneTex, catalog 43019), anti-E-Cadherin (E-cad, 1:200, Proteintech, catalog 60335), anti-Pax6 (1:200, Proteintech, catalog 12323) and anti-Ki67 (1:1000, Cell signaling technology, catalog 9449) at 4 °C overnight. The primary antibodies were then washed off and sections were incubated with secondary antibodies (Invitrogen, America) for 1 h at room temperature. Cells were counterstained with Hoechst for 10 min after wash. Images were captured using a spinning disk confocal microscope (Andor Technology, UK).

Stroke leads to damage in the cerebral cortex, the atrophy was more severe without treatment, therefore we chose to observe the neuron recovery corresponding to the atrophied area in the model group. For staining of mice brain tissues, mice were anesthetized and immediately perfused with PBS followed by 10% formalin for 30 min. Brains were fixed overnight in fixative at 4 °C. Fixed brains were dehydrated in 30% sucrose in PBS for 2 days at 4 °C. Brains were embedded in the OCT compound (SAKURA, Japan). Brain sections were obtained at a thickness of 25 µm using a microtome cryostat (Leica, Germany). Tissues were permeabilized, blocked and incubated as the above-mentioned protocol for cultured cell staining. Tissues were incubated with primary antibodies, including anti-NeuN (1:300, Abcam, catalog ab177487), anti-GFAP (1:1000, Cell signaling technology, catalog 3670S), anti-ß-III- tubulin (1:300, Cell signaling technology, catalog 5568S) or anti-Nestin (1:250, Santa Cruz Biotechnology, catalog Sc-23927). For TUNEL staining, in situ cell death detection kit (Roche, Germany) was used to detect the cell apoptosis according to the manufacturer’s instructions. Briefly, 3% BSA incubated sections were incubated with TUNEL reaction mixture for 1h at 37 °C in the dark. Then sections were incubated with anti-NeuN primary antibody (1:300, Abcam, catalog ab177487) and corresponding secondary antibody successively.

### Western blot analysis

Total protein was extracted from NSCs or exosomes using RIPA lysis buffer (Biosharp, China) with protease inhibitor PMSF. Protein content was observed by the BCA assay (Biosharp, China). Protein samples (30 µg) were electrophoretically separated on 12% SDS-PAGE gels and then transferred to polyvinylidene fluoride membranes (PVDF, Immobilon, America). The membranes were incubated with primary antibodies including TSG101 (1:1000, ABclonal, catalog A1692), CD63 (1:1000, ABclonal, catalog A19023) and CD9 (1:2000, ABclonal, catalog A19027) overnight at 4 °C. The membranes were next incubated with secondary antibodies for 1 h at room temperature (1:5000, Proteintech) and were detected using the ECL Enhanced Kit (ECL, Abclonal).

### Motor function assessment

Testing on balance beam, ladder rung, rotarod test and mNSS tasks was conducted preoperatively, and at 1 to 8 weeks postoperatively. Investigators were blinded to treatment groups in test.

**Balance beam:** The balance beam apparatus used in this study was a 10 mm square wood in width and 50 cm wood in length (Beijing Zhongshi Science, China). Mice were trained to pass through the balance beam 3 days before the MCAO/R procedure. The mice that successfully passed the beam without foot slips were recruited and grouped.

On behavioral test days (0, 4, 8 weeks after treatment), the right feet slips were recorded when mice were passing through the balance beam three times. The scores were full score (10) minus the number of foot slips. If the mouse could not pass through or fall, the minimum score was recorded as 0.

**Ladder rung:** The ladder rung instrument was made up of two transparent glass walls and 70 irregular metal bars (Beijing Cinontech Co. Ltd, China). The number of mice that stepped wrong was recorded on behavioral test days. The score was calculated as 10 minus the number of wrong steps. If the mouse could not pass through, the minimum score was recorded as 0.

**Rotarod test:** At 3 days before MCAO/R procedure, mice were trained on an accelerating rotarod at 30 rpm and only the mice that remained on the rotarod for 300 s at 30 rpm of three trials were recruited and grouped. The test was carried out at 30 rpm on behavioral days. The final scores were the seconds of mice remaining on the rotarod over three trials. The maximum score is 300 seconds.

**mNss:** According to the aforementioned report (Supplementary table 1 (Chen *et al.,* 2001)), mNss test is a composite of balance, motor, and reflex tests to assess neurological deficit. The normal score varies from 0 to 10, where 0 represents normal function and 10 maximal deficits. Three measurements were obtained per behavioral day.

### Golgi staining and analysis

Golgi staining was conducted using the FD rapid Golgi Stain kit (FD Neuro Technologies, America) according to the manufacturer’s instructions. Mice were anesthetized and sacrificed, and the brains were removed quickly and immersed in the mixture of Solution A and B for 2 weeks at room temperature in the dark. The brain was then transferred into Solution C for 48 h at 4 °C. Sections were cut with 100 µm thickness using a concussion slicer (Lecia, Germany) and stained with D and E mixture. Images were captured by an inverted microscope using Z-stack images (Lecia, Germany). Golgi-stained neurons were reconstructed using Fiji-Image J. The total dendritic length, the number of dendritic spines and intersections were calculated and analyzed by Sholl analysis according to the previous study (Yang *et al.,* 2020).

### Nissl staining

Nissl staining was conducted using the Nissl Stain Solution (Solarbio, America) according to the manufacturer’s instructions. Mice brain sections were stained with methylene blue stain for 10 min at 65°C, then differentiated by nissl differentiation solution for 3 min. The brain sections were subsequently treated in ammonium molybdate solution for 5 min followed by a quick rinse quickly in distilled water to avoid decolorizing. Images were taken using an inverted microscope (Lecia, Germany).

### Oxygen-glucose deprivation and reoxygenation (OGD/R)

To perform OGD/R on cultured NSCs, the normal culture medium was replaced with Dulbecco’s modified of eagle’s medium (Solarbio, China). The culture was then incubated in a hypoxia chamber aerated with 5% CO_2_, 94% N_2_ and 1% O_2_ at 37° C for 2 h. Then the NSCs were transferred back into the normal culture medium and incubated in normal culture conditions for 24 h.

### Intracellular ROS detection

ROS level was detected using the fluorescent probe DCF-DA (Invitrogen, America). Cultured NSCs were incubated with 10 µM DCF-DA for 30 minutes at 37 °C and then fixed with 4% paraformaldehyde. DCF fluorescence was photographed and quantified via a spinning disk confocal microscope (Andor Technology, UK).

### MDA level measurement

The MDA level was measured at 3 days after treatment by the TBA method (Nanjing Jiancheng Bioengineering Institute, China) according to the manufacturer’s instructions. The ipsilateral brain was homogenized and incubated in the assay solution at 95°C for 80 min. The optical density was measured at 532 nm by a microplate reader.

The value was calculated based on the standard formula.

### Microarray analysis of exosomal miRNAs

Sequencing libraries of miRNAs of NSC-derived exosomes were produced using NEBNext Multiplex Small RNA Library Prep Set for Illumina (NEB, United States) following the previous report (Cai *et al.,* 2021).

The downstream target genes of exosomal miRNAs were predicted using three online databases: Targetscan (http://www.targetscan.org/), miRcode (http://www.mircode.org/) and miRDB (http://mirdb.org/). The enrichment analysis of the predicted target genes was performed using ClusterProfiler R package for GO process and KEGG pathway enrichment. The miRNA-mRNA regulatory network was built by Cytoscape software. miRNA was calculated by qPCR. Libraries were prepared by ligating adaptors to the total RNA, PCR amplification and size selection using 6% polyacrylamide gels. Sequencing was performed on Illumina NovaSeq 6000 (Illumina Inc, USA).

### Statistics

GraphPad Prism version 7 was used for statistical analyses. Unpaired t-tests (two-tailed) were used for single comparisons, and two-way ANOVA was used for multiple comparisons. Survival analysis was performed via the Kaplan-Maier method. All data are presented as mean ± SEM.

## Acknowledgments

This work is supported by National Natural Science Foundation of China (Grant No. 32070973, 31871481), Fundamental Research Funds for the Central Universities (Program No. 2662022JC002).

## Additional information

## Funding

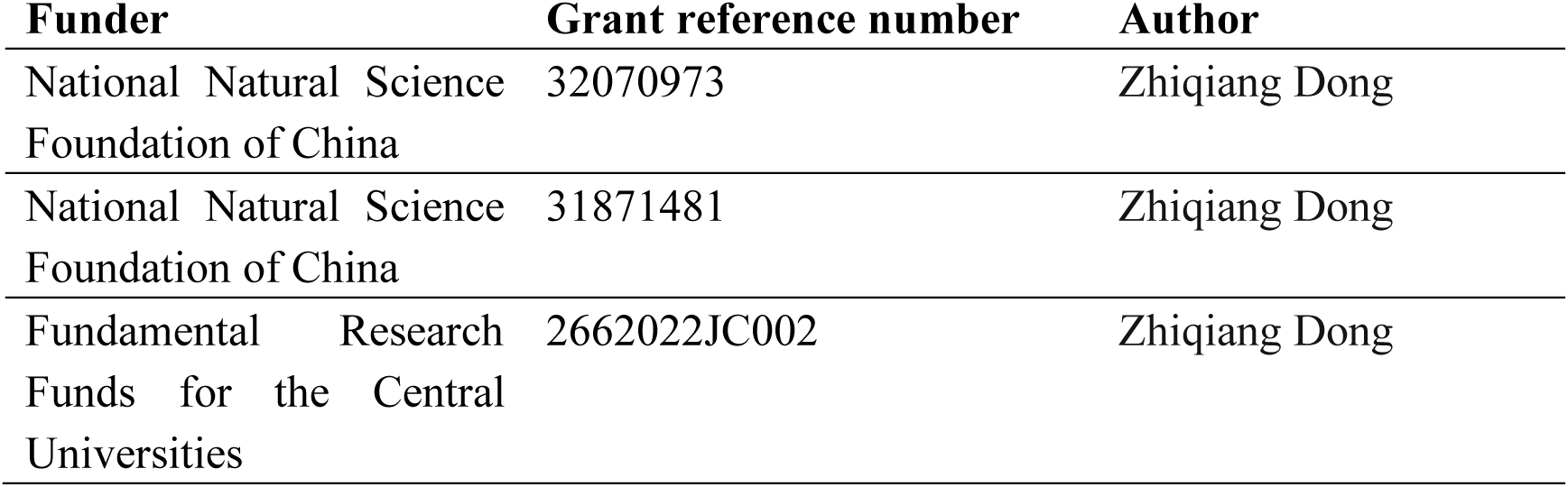

### Author contributions

Ruolin Zhang, Conceptualization, Methodology, Software, Validation, Formal analysis, Investigation, Resources, Data curation, Writing - original draft preparation, Writing - review & editing; Visualization; Weibing Mao, Methodology, Validation; Lumeng Niu, Methodology, Validation; Wendai Bao, Conceptualization, Writing - review & editing; Yiqi wang, Software, Visualization; Zhihao Yang, Methodology, Investigation; Yasha Zhu, Methodology; Haikun Song, Methodology, Investigation; Jincao Chen, Conceptualization,; Guangqiang Li, Methodology; Meng Cai, Project administration, Writing - review & editing; Zilong Yuan, Validation; Jiawen Dong, Validation; Min Zhang, Conceptualization; Nanxiang Xiong, Project administration, Writing - review & editing; Jun Wei, Conceptualization, Project administration, Writing - original draft, Supervision; Zhiqiang Dong, Conceptualization, Project administration, Writing - review & editing, Writing - original draft, Supervision, Funding acquisition

### Ethics

All animal procedures were performed in compliance with guidelines for the care and use of animals and were approved by the University Huazhong Agriculture Institutional Animal Care and Use Committee (Approval number: HZAUMO-2021-0111).

### Data and materials availability

All data are available in the main text or the supplementary materials.

---

**Figure 1 - supplement 1.**
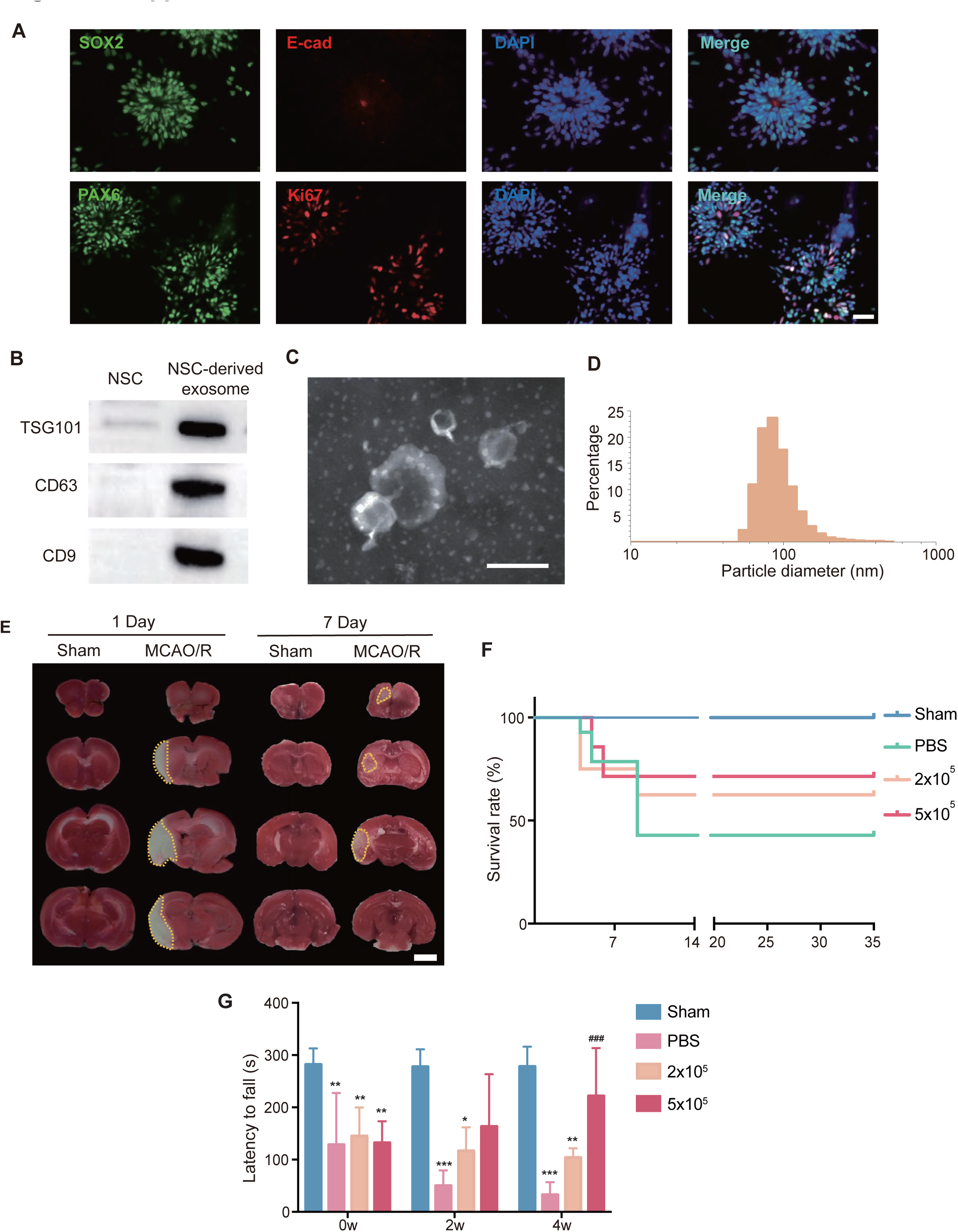
Characterization of NSCs and NSC-derived exosomes, and effects of different doses of transplanted NSCs. (A) Representative images of immunostaining against SOX2 (green), E-cad (red), PAX 6 (green), Ki67 (red), and DAPI (blue) on NSCs. Scale bar: 25 µm. (B) Western blot results showed the expression of exosome markers TSG101, CD63 and CD9. (C) Representative TEM image of NSC-derived exosomes. Scale bar: 200 nm. (D) The distribution of particle diameters of exosomes mixture examined by Malvern Nano ZS90. \*\*\**P*<0.001. (E) Representative images of TTC staining on day 1 and day 7 post-stroke. The infract areas were marked by dotted lines. Scale bar: 2 mm. (F) The survival curve of different dosed NSC treatment (2×10^5^ and 5×10^5^). (G) The rotarod test at 2 weeks and 4 weeks after treatment, n = 5 mice / group. \**P* < 0.05, \*\**P* < 0.01, \*\*\**P* < 0.001 versus Sham group. ^###^ *P* < 0.001 versus Model group. ns indicates non-significant difference. Figure 1 **- supplement 1-source data 1.** Characterization of NSCs and NSC-derived exosomes, and effects of different doses of transplanted NSCs. Figure 1 **- supplement 1-source data 2.** Characterization of NSCs and NSC-derived exosomes, and effects of different doses of transplanted NSCs.

**Figure 1 - supplement 2.**
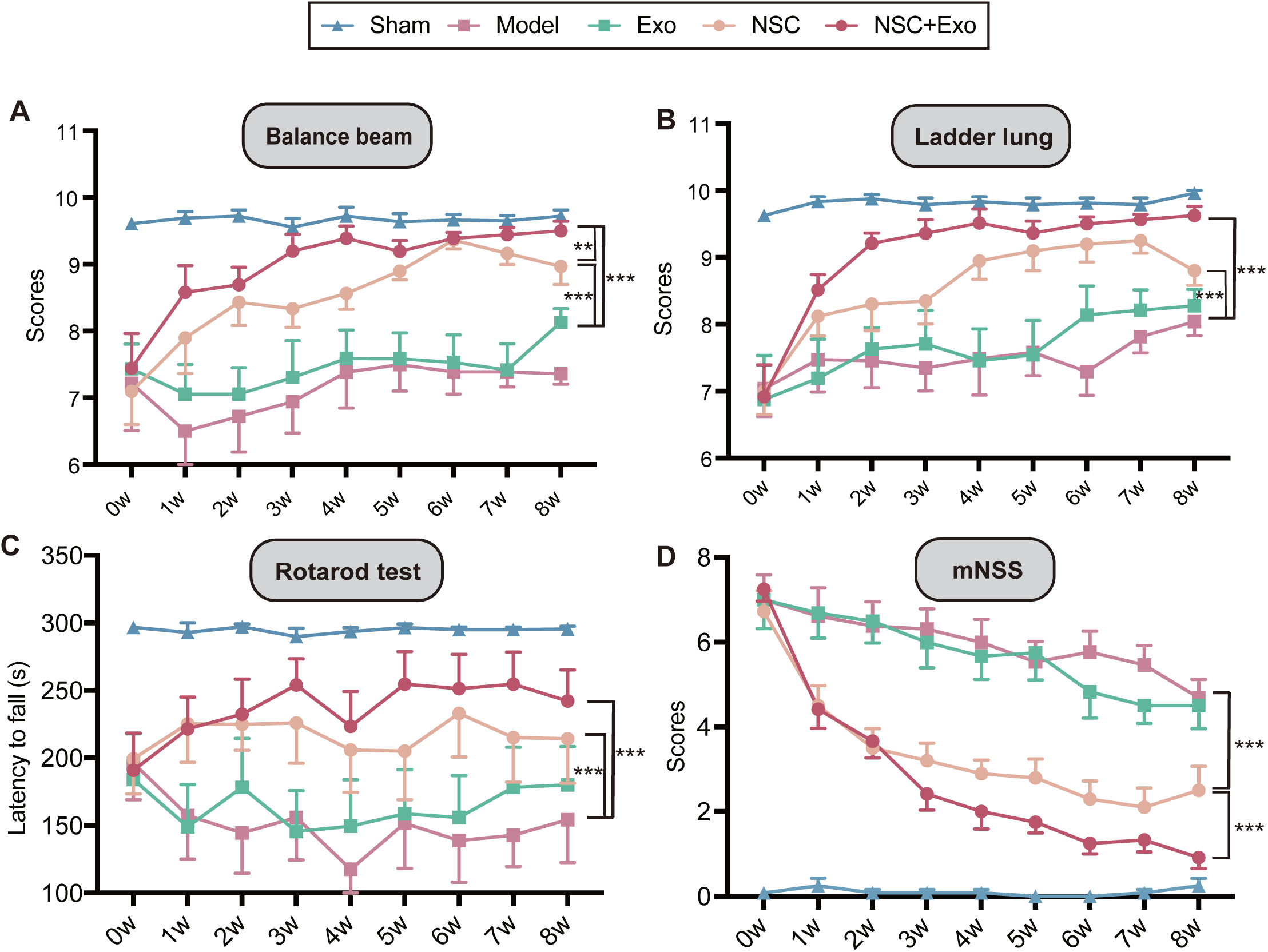
Effects of different treatment strategies of NSCs and exosomes on ischemic stroke in MCAO/R mice. Behavioral test results (the balance beam (A), ladder rung (B), rotarod tests (C)) and mNSS (D) up to 8 weeks after treatment, n = 10 mice per group. * *P* < 0.05, ** *P* < 0.01, *** *P* < 0.001. Figure 1 **- supplement 2-source data 1.** Effects of different treatment strategies of NSCs and exosomes on ischemic stroke in MCAO/R mice.

**Figure 2 - supplement 1.**
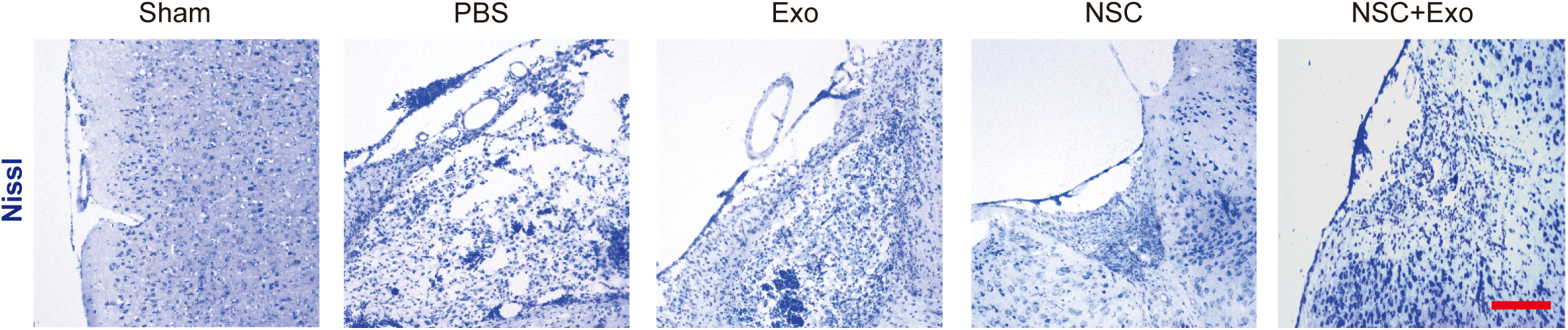
NSCs and Exosomes combination treatment reduced the neuronal loss in the ipsilesional hemisphere. Nissl staining of infarct area in the brain at 4 weeks after treatment. Scale bar: 25 µm.

**Figure 3 - supplement 1.**
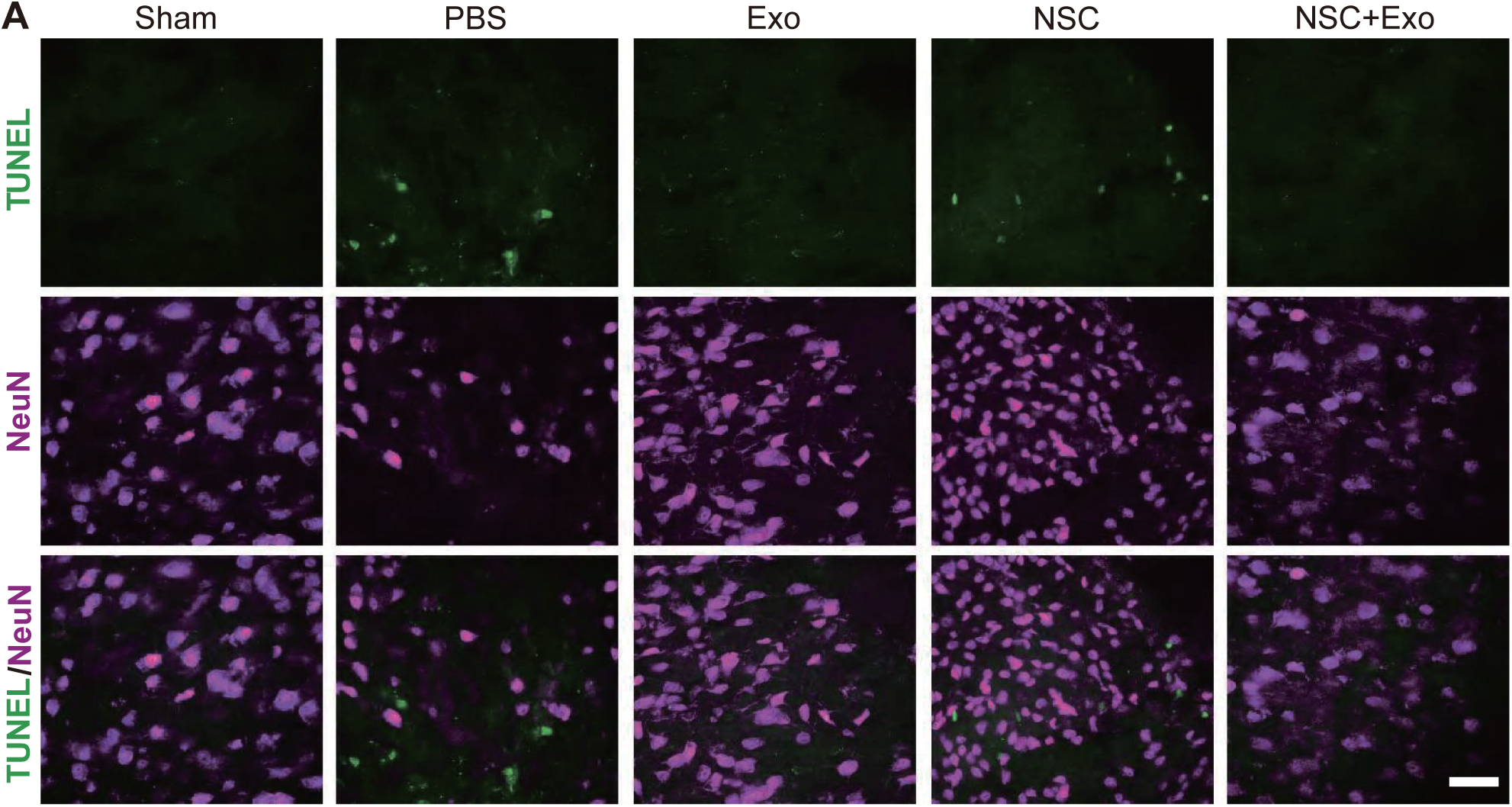
Exosome transplantation reduced the excessive apoptosis cells after MCAO/R. TUNEL (green) and NeuN (purple) staining at 7 days after treatment. Scale bar: 50 µm.

**Figure 4 - supplement 1.**
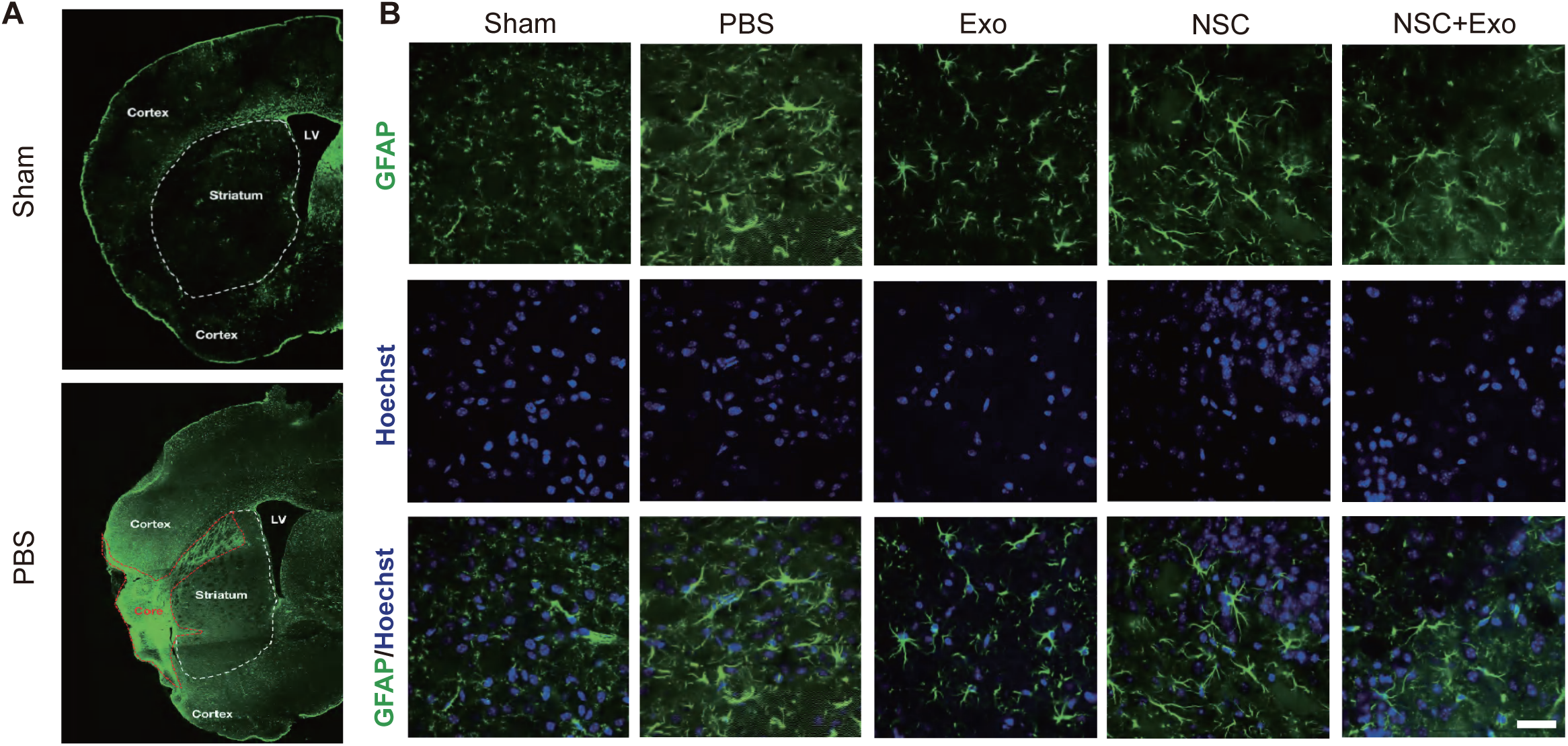
Combined treatment of NSCs and exosomes reduced the formation of glial scars in MCAO/R mice. (A) Representative images of GFAP staining at 8 weeks after treatment. Scale bar: 2 mm. (B) Representative images of GFAP staining at 7 days after treatment. Scale bar: 50 µm.

**Figure 5 - supplement 1.**
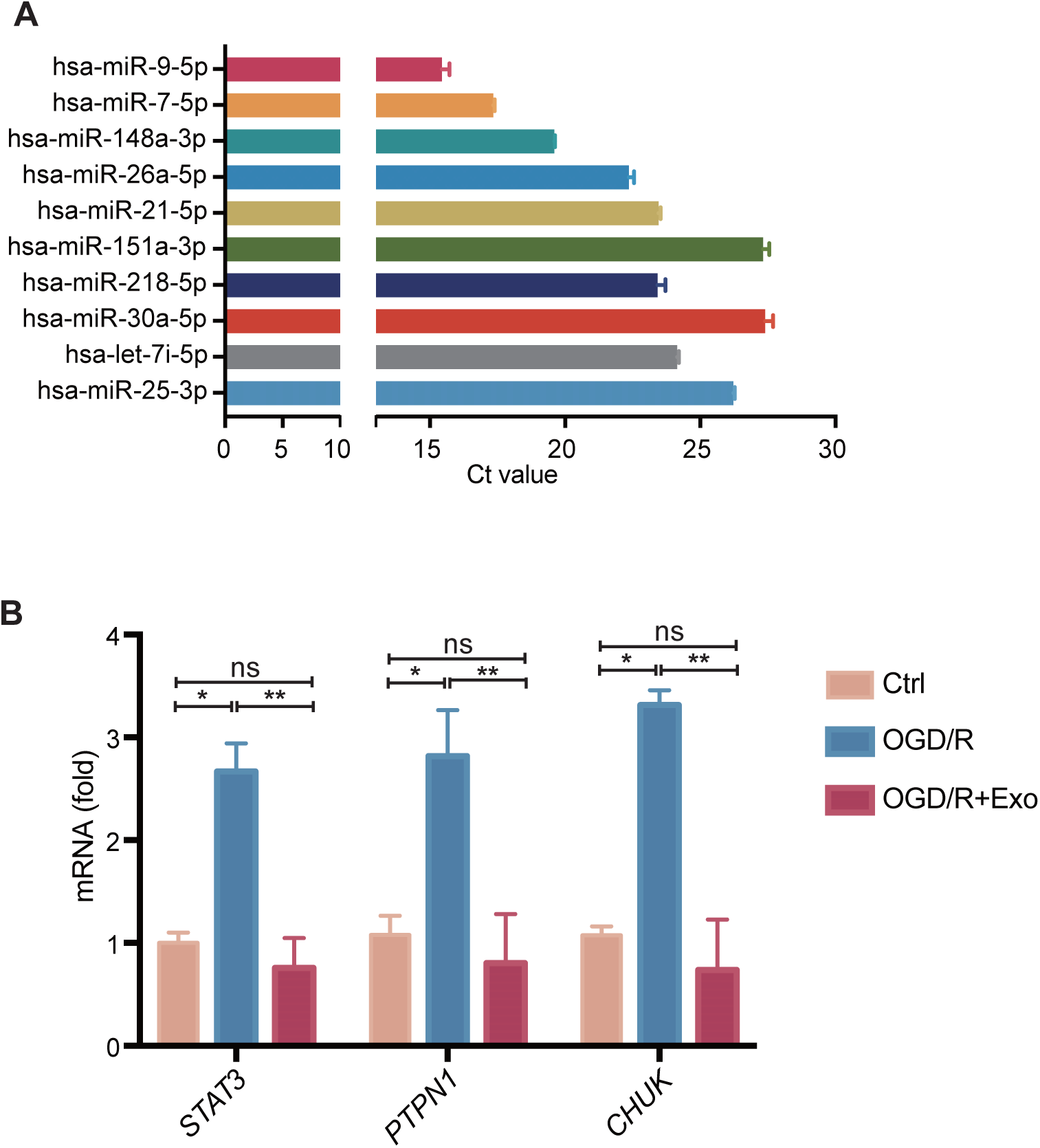
Verification of top 10 abundant miRNAs and the expression of the selected target genes after OGD/R. (A) Cycle threshold (Ct) value of each top 10 abundant miRNA detected by qPCR. (B) The relative mRNA expression level of *STAT3, PTPN1* and *CHUK* after OGD/R treatment. * *P* < 0.05, ** *P* < 0.01. ns indicates non-significant difference. Figure 5 **- supplement 1-source data 1.** Verification of top 10 abundant miRNAs and the expression of the selected target genes after OGD/R.

